# Engineering human Mucosal Associated Invariant T (MAIT) cells with chimeric antigen receptors for cancer immunotherapy^1^

**DOI:** 10.1101/2022.07.28.501764

**Authors:** Mikail Dogan, Ece Karhan, Lina Kozhaya, Lindsey Placek, Xin Chen, Mesut Yigit, Derya Unutmaz

## Abstract

Engineering immune cells with chimeric antigen receptors (CAR) is a promising technology in cancer immunotherapy. Besides classical cytotoxic CD8+ T cells, innate cell types such as NK cells have also been used to generate CAR-T or CAR-NK cells. Here we devised an approach to program a non-classical cytotoxic T cell subset called Mucosal Associated Invariant T (MAIT) cells into effective CAR-T cells against B cell lymphoma and breast cancer cells. Accordingly, we expressed anti-CD19 and anti-Her2 CARs in activated primary human MAIT cells and CD8+ T cells, expanded them *in vitro* and compared their cytotoxicity against tumor cell targets. We show upon activation through CARs, CAR-MAIT cells exhibit high levels of cytotoxicity towards target cells, comparable to CD8+ CAR-T cells, but interestingly expressed lower levels of IFN-γ than conventional CAR CD8+ T cells. Additionally, in the presence of vitamin B2 metabolite 5-ARU, which is a conserved compound that activates MAIT cells through MHC-I related (MR1) protein, MAIT cells killed MR1-expressing target breast cancer and B cell lymphoma cell lines in a dose dependent manner. Thus, MAIT cells can be genetically edited as CAR-T cells or mobilized and expanded by MR1 ligands as an off-the-shelf novel approach to cell-based cancer immunotherapy strategies <u>while being comparable to conventional methods in effectivity</u>.

**Key Points:**

- MAIT cells expressing CARs effectively kill CD19 and Her2 expressing targets
- CAR-MAITs display strong cytotoxicity with lower TNF-γ compared to CD8+ CAR-T cells

## Introduction

Chimeric antigen receptor (CAR) based approaches have been one of the most promising immunotherapies in the last decade. The majority of CAR immunotherapies are focused on the engineered conventional CD8+ T cells against tumor targets, referred to as CAR-T cells.(1) While other cell types such as natural killer cells,(2) γδT cells(3) and macrophages(4) have also been tested as CAR-modified cells, different subtypes of T cell family and the innate cells remain to be tested to determine their effectiveness, durability and safety for CAR immunotherapies.

Human Mucosa associated invariant T (MAIT) cells constitute a large subset of innate-like T cells that recognize conserved bacterial ligands from vitamin B metabolic pathways (B_2_, Riboflavin and B_9_, Folate). (5-7) Predominantly residing in tissue, MAIT cells account for 1 - 8% of peripheral T lymphocytes and up to 40% of tissue specific T cells.(8-10) An invariant T cell receptor (TCR) alpha chain (Vα7.2-Jα33) integrated with a narrow TCR β-chain repertoire is used to identify MAIT cells.(10) MAIT cells recognize their specific antigens through major histocompatibility complex class I-related protein (MR1) which potentially can allow them to avoid attacking the host cells during a cell-based immunotherapy, contrary to conventional T cells, which use polymorphic MHC molecules for antigen recognition. In addition to Vα7.2 TCR, MAIT cells are also characterized by a high expression of CD161 and IL18R surface markers, and thus can consequently be identified as TCR Vα7.2+CD161+IL-18Rα+ αβ T cells.(10, 11)

MAIT cells are activated through vitamin B2 metabolites presented on MR1 molecules on antigen presenting cells. Upon activation, they secrete cytokines such as TNF-γ, TNF-α and IL-17 and display cytotoxicity through perforin and granzyme expression(12-15). Several studies have reported that MAIT cells are abundant in the tumor microenvironment(16-18). Their tendency to naturally migrate to the tumor microenvironment can be exploited for immunotherapy, such as CAR engineering approaches.

Here we engineered MAIT cells with chimeric antigen receptors using CD19 and Her2 expressing cell lines as targets. MAIT cells expressing anti-Her2 or anti-CD19 (herein named CAR-MAIT cells), showed high cytotoxicity to target cells and expressed low levels of pro-inflammatory cytokines such as TNF-γ. Additionally, CAR-MAIT cells were able to kill their targets in the presence of vitamin B2 metabolites in a dose dependent manner, which can be a method to mobilize them towards MR1 expressing tumor cells. Taken together, these findings suggest that MAIT cells can be novel candidate cell types for cancer immunotherapies given their cytotoxic abilities that are comparable to conventional CAR-T cells, combined with their “off-the-shelf” generation potential.

## Methods

### PBMC and T cell purification

Healthy adult blood was obtained from AllCells® Quincy, MA. PBMCs were isolated using Ficoll-paque plus (GE Health care). CD4+T, CD8+ T, CD19+ B cells, CD14+ monocytes were purified using Dynal CD4 Positive, CD8 Positive, CD19 Positive Isolation Kits (all from Invitrogen) and CD14 Mircrobeads Ultra-pure (Miltenyi Biotec). To purify Vα7.2 positive cells, CD4 Negative fraction of PBMCs was used. Cells were stained with Vα7.2 PE conjugated antibody for 30 minutes at 4°C, washed with PBS (Phosphate Buffered Saline) containing 5% FBS (Fetal Bovine Serum) then labeled with anti-PE microbeads (Miltenyi Biotec Inc.) for 15 minutes at 4°C. After washing, labeled cells were resuspended in PBS+5% FBS, passed through cell strainer then separated on autoMACS Pro Separator for isolation of Vα7.2 positive and negative cells. CD4+ T, CD8+ T, CD 19+ B cells and CD 14+ monocytes were >99% pure and assessed by flow cytometry staining with respective antibodies. Vα7.2+ T cells were >95% pure and verified by flow cytometry. Verification of sorted MAIT cells is shown in Supplementary Figure 4.

### Designing CAR and MR1 overexpression constructs

CAR constructs consist of CD8 alpha signal peptide, single chain variable fragment (scFv) of a CD19 or Her2 antibody, CD8 hinge domain, CD8 transmembrane domain, 4-1BB (CD137) intracellular domain and CD3ζ domain which were designed with Snapgene and synthesized via Genscript. CD8a signal peptide, CD8 hinge, CD8 transmembrane domain, 4-1 BB intracellular domain and CD3ζ domain sequences were obtained from Ensembl Gene Browser and codon optimized with SnapGene by removing the restriction enzyme recognition sites that are necessary for subsequent molecular cloning steps while preserving the amino acid sequences. Anti-CD19 and anti-Her2 scFv amino acid sequences were obtained from Addgene plasmids #79125 and #85424, respectively, reverse translated to DNA sequences and codon optimized with Snapgene 5.2.4. Human MR1 transcript variant 1 (NM_001531) cDNA clone was purchased from Origene. The constructs were then cloned into a lentiviral expression vector with a multiple cloning site separated from GFP reporter via an Internal Ribosomal Entry Site (IRES).

### Lentiviral production and titration

Cloned lentiviral constructs including anti-CD19 CAR, anti-Her2 CAR and MR1 encoding vectors were co-transfected with the packaging plasmids VSVG, pLP1 and pLP2 into 293 cells using Lipofectamine™ 3000 (Invitrogen) according to the manufacturer’s protocol. Viral supernatants were collected 24-48 hours post-transfection, filtered through a 0.45 μm syringe filter (Millipore) to remove cellular debris, and concentrated with Lenti-X (Invitrogen) according to the manufacturer’s protocol. Lentivirus supernatant stocks were aliquoted and stored at −80°C. To measure viral titers, virus preps were serially diluted on Jurkat cells. 72 hours after infection, GFP positive cells were counted using flow cytometry and the number of cells transduced with virus supernatant was calculated as infectious units per ml. The cells were cultured in complete RPMI 1640 medium (RPMI 1640 supplemented with 10% FBS; Atlanta Biologicals, Lawrenceville, GA), 8% GlutaMAX (Life Technologies), 8% sodium pyruvate, 8% MEM vitamins, 8% MEM nonessential amino acid, and 1% penicillin/streptomycin (all from Corning Cellgro) for 72 hours (19). %0.05 Trypsin-0.53 mM EDTA (Corning Cellgro) was used to detach adherent MDA-231 cells.

### Engineering CAR-T cells and target tumor cell lines

CD8+ T cells were activated using anti-CD3/CD28 coated beads (Invitrogen) at a 1:2 ratio (beads:cells) and infected with anti-CD19 CAR, anti-Her2 CAR or empty lentivectors with multiplicity of infection (MOI) of 3. Cells were then expanded in complete RPMI 1640 medium supplemented with 10% Fetal Bovine Serum (FBS, Atlanta Biologicals), 1% penicillin/streptomycin (Corning Cellgro) and 10ng/ml of IL-2 for 10-12 days and cultured at 37°C and 5% CO2 supplemented incubators. For PBMC assays, cells were activated with 30μM 5-amino-4-D-ribitylaminouracil dihydrochloride (5-ARU, Toronto Research Chemicals, Canada). IL-2 was added 72 hours after their activation in order to avoid non-MAIT cells growth. Respective viruses were added 24 hours after the activation. Cells were expanded for 10-12 days and cytotoxicity assays were performed following their expansion. In Figure 1b, T2 cells were added to PBMC cultures on day 8 post activation then cocultures were analyzed by flow cytometry 3 days later. To generated and expand pure primary MAIT cells, sorted Vα7.2+ cells were cultured overnight in complete medium containing IL-7+IL-15 (20ng/ml). At the second day, cells were activated with 30μM 5-ARU in the presence of autologous CD 14+ monocytes and CD 19+ primary B cells. 20ng/ml IL-2 was added the next day along with either anti-CD19 CAR lentivirus, anti-Her2 CAR lentivirus or empty vector as a control. A portion of the activated MAIT cells were left non-infected and expanded for 3 weeks and used in Figure 3f. For MR1 overexpressing MDA-231, wild-type MDA-231 cells were transduced with 3 MOI of MR1 overexpression lentivirus and proliferated. The infection levels were determined by GFP expression on Flow Cytometry analysis.

**Figure 1:**
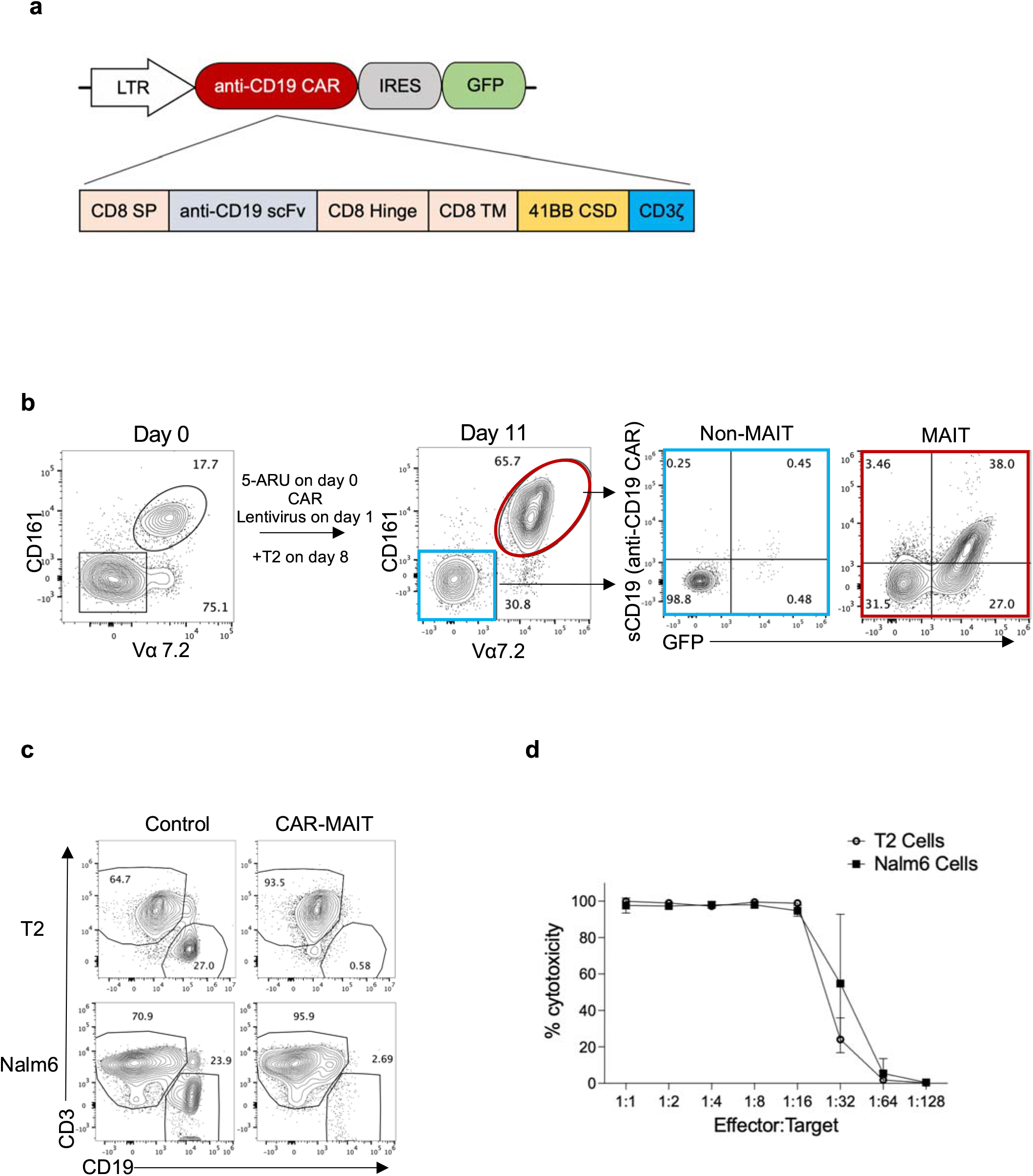
Engineering human primary MAIT cells into CAR-MAIT cells. **a**. Schematic illustration of anti-CD19 CAR construct. A constitutive LTR promoter drives the anti-CD19 CAR and GFP genes that separated by an Internal Ribosomal Entry Site (IRES). Anti-CD19 CAR construct consists of CD8 alpha signal peptide, single chain variable fragment of an anti-CD19 antibody, CD8 Hinge, CD8 transmembrane domain, 4-1BB (CD137) co-stimulatory domain and CD3ζ domain, b. Flow cytometry plots demonstrating the proliferation and transduction of primary MAIT cells with an anti-CD19 CAR construct. Vα7.2+CD161+ CD8+ T cells within PBMC were activated with the riboflavin metabolite, 5-amino-6-D-ribitylaminouracil (5-ARU), transduced with a lentivirus encoding CD 19 targeting chimeric antigen receptor (CAR) and GFP marker, then target T2 cells were added on day 8 post activation and cultures were analyzed by flow cytometry 3 days later. Non-MAIT T cells are shown in blue boxes and MAIT cells are indicated with a red circle and a red box. c. Representative flow cytometry plots of CAR-MAIT cytotoxicity assays with CD19+ T2 and Nalm6 target cells. MAIT cells transduced with an empty lentivector were used as controls. Effector MAIT cells were identified with CD3 staining while target cells were shown via CD19 staining. Effector (CAR expressing T cells) to target ratio of 1:1 is shown, d. Line graph representing CAR-MAIT cytotoxicity against Nalm6 and T2 cells. Circles and squares indicate T2 and Nalm6 cell cytotoxicity curves, respectively. Error bars represent one standard deviation of mean value. The effector to target cell ratio was calculated based on the number of effector cells expressing CARs. Effector to target ratio was titrated from 1:1 to 1:128 with 2-fold reciprocal dilutions of effector cells. Percent cytotoxicity was calculated based on the percentage of cell death in experimental conditions in relation to control conditions. Each experiment was performed three times.

### Staining and flow cytometry analysis

Cells were resuspended in staining buffer (PBS + 2% FBS) and incubated with fluorochrome-conjugated antibodies for 30 min at 4°C. Identification of MAIT T cells was determined by staining with Vα7.2-PE and CD161-BV421 antibodies (Biolegend). CD4^-^ population for calculating MAIT cell percentage in PBMC was determined by gating out CD4+ cells stained with CD4-BV605 antibody (Biolegend). CAR CD8+ T cells were identified with CD3-APC/Cy7 or CD8-Alexa 700 antibodies (Biolegend). Activation of CAR-MAIT and CAR CD8+ T cells were determined with CD25 staining using CD25-APC antibody (Biolegend). CAR expression was determined with Human Her2 / ErbB2 Protein, Fc Tag (Acro) or Human CD19 (20-291) Protein, Fc Tag, low endotoxin (Super affinity) (Acro) followed by a secondary staining with APC conjugated anti-human IgG Fc Antibody (Biolegend). For cytotoxicity assay analysis, T2, Nalm6 and primary B cells were labeled with PE/Cy7 anti-human CD19 Antibody (Biolegend) and MDA-231 cells were stained with APC conjugated anti-human CD340 (erbB2/HER-2) Antibody (Biolegend). MR1 expression on T2 cells was assessed with anti-MR1-PE (Biolegend). Samples were acquired on a BD *FACSymphony A5 analyzer* and data were analyzed using FlowJo (BD Biosciences).

### Cytotoxicity assays

Following the expansion of effector cells for 10-12 days, the cells were analyzed for their GFP and CAR expressions. Effector cell to target cell ratio was calculated based on the number of CAR expressing cells. CAR expressing cells were titrated from 1:1 to 1:128 with 2-fold dilutions while the target cell number was constant. For primary MAIT cell cytotoxicity assay, 5-ARU was titrated from 30 μM to 0.003 μM with 3-fold reciprocal dilutions. Cytotoxicity assay conditions were analyzed with Flow Cytometry at 72 hours of co-incubation. The cells were first gated based on their forward and side scatter properties to exclude doublets, debris and dead cells. Viability dye staining was included in the original experiments (Fig 1 d) and showed that forward and side scatter gating was excluding dead and dying cells. Target cells were identified with CD19 (primary B cells, T2 and Nalm6 Lymphoma cell lines) and Her2 staining (MDA-231, Breast cancer cell line) and effector cells were identified with CD3 staining and GFP expression from CAR-GFP lentivector. Cytotoxicity for each Effector:Target ratio condition was calculated based on the corresponding control condition with the following formula: (Percentage of target cells in control condition - percentage of target cells in experimental condition) / percentage of target cells in control condition * 100.

### Cytokine Assay

Cell culture supernatants from cytotoxicity assays were collected 72 hours after combining the effector and the target cells and were immediately stored in -80°C. Secreted proteins and cytokines, including TNF-γ, TNF-α, GM-CSF and Granzyme B production was measured using Qbeads Plex screen assay (Essen BioScience, Inc, USA) according to the manufacturer’s instructions and analyzed using iQue Screener Plus (IntelliCyt, USA).

### Statistical Analyses and Reproducibility

All statistical analyses were performed using GraphPad Prism V8 software. Two-tailed unpaired t test was used to determine the statistical significances and exact P values are reported. P < 0.05 was considered significant. Numbers of repeats for each experiment were described in the associated figure legends.

## Results

### Activation and cytotoxicity of anti-CD19 CAR engineered MAIT cells

In order to engineer primary MAIT cells into CAR-MAITs, we first designed a lentiviral CAR construct composed of a CD8α signal peptide, an anti-CD19 antibody single chain variable fragment (scFv), CD8 hinge domain, CD8 transmembrane domain, 41BB co-stimulatory domain and CD3 zeta (ζ) domain. The CAR construct was designed to be co-expressed with a GFP marker via an internal ribosome entry site (IRES) under LTR promoter (Fig. 1a). To express the CAR construct and expand MAIT cells, we first activated MAIT cells by stimulating these cells in PBMC with 5-amino-6-D-ribitylaminouracil (5-ARU), a vitamin B2 biosynthesis derivative. Within the PBMC, MAIT cells were defined as CD4^-^ or CD8+ T cells expressing both TCR Vα7.2 and CD161 surface markers(20)(Fig. 1b). One day after stimulating MAIT cells in PBMC with 5-ARU, cells were transduced with lentivirus encoding the CAR construct. The cells were cultured for 7 days in the presence of IL-2 then T2 cells were added at 1:1 effector to target cell ratio. Three days later cocultures were stained for CD8, CD161, TCR Vα7.2 and anti-CD19 CAR expression. After 11 days, activated MAIT cells were selectively expanded and therefore constituted the majority of CD8+ T cells (Fig. 1b, Supplementary Fig. 1a). MAIT cells also selectively expressed anti-CD19 CAR compared to Vα7.2 CD16T non-MAIT cells, corresponding to the marker GFP levels (Fig. 1b).

After confirming CAR expression in MAIT cells, we tested their cytotoxic function using T2 and Nalm6 cell lines as target cells known to express CD 19 on their surface (Supplementary Fig. 1b). Flow cytometry analysis 72 hours after co-culture revealed that anti-CD19 CAR-MAIT cells were able to kill their targets (Fig 1c). A more detailed cytotoxic assay testing different effector:target ratios showed that MAIT cells were highly cytotoxic, killing target cells up to an effector:target ratio of 1:32 (Fig 1d). Together, these findings demonstrate the specific killing of target cells by CAR-MAIT cells.

### Expansion, isolation and cytotoxicity of purified CAR-MAIT cells to target cancer cell lines

To further investigate the possibility of using MAIT cells as candidates for cancer immunotherapies, we developed a sorting and expansion strategy to generate purified CAR-MAIT cells for further analyses (Fig. 2a). Given that the majority of MAIT cells do not express CD4(21), we used the CD4 negative fraction of PBMC to sort TCR Vα7.2+ cells. MAIT-enriched Vα7.2+ cells were then activated with 5-ARU in the presence of autologous CD19+ B cells and CD14+ monocytes as antigen presenting cells (APCs) and transduced with CAR-expressing lentiviruses. Cells were then expanded in the presence of IL-2 for 10-12 days (Fig. 2a).

**Figure 2:**
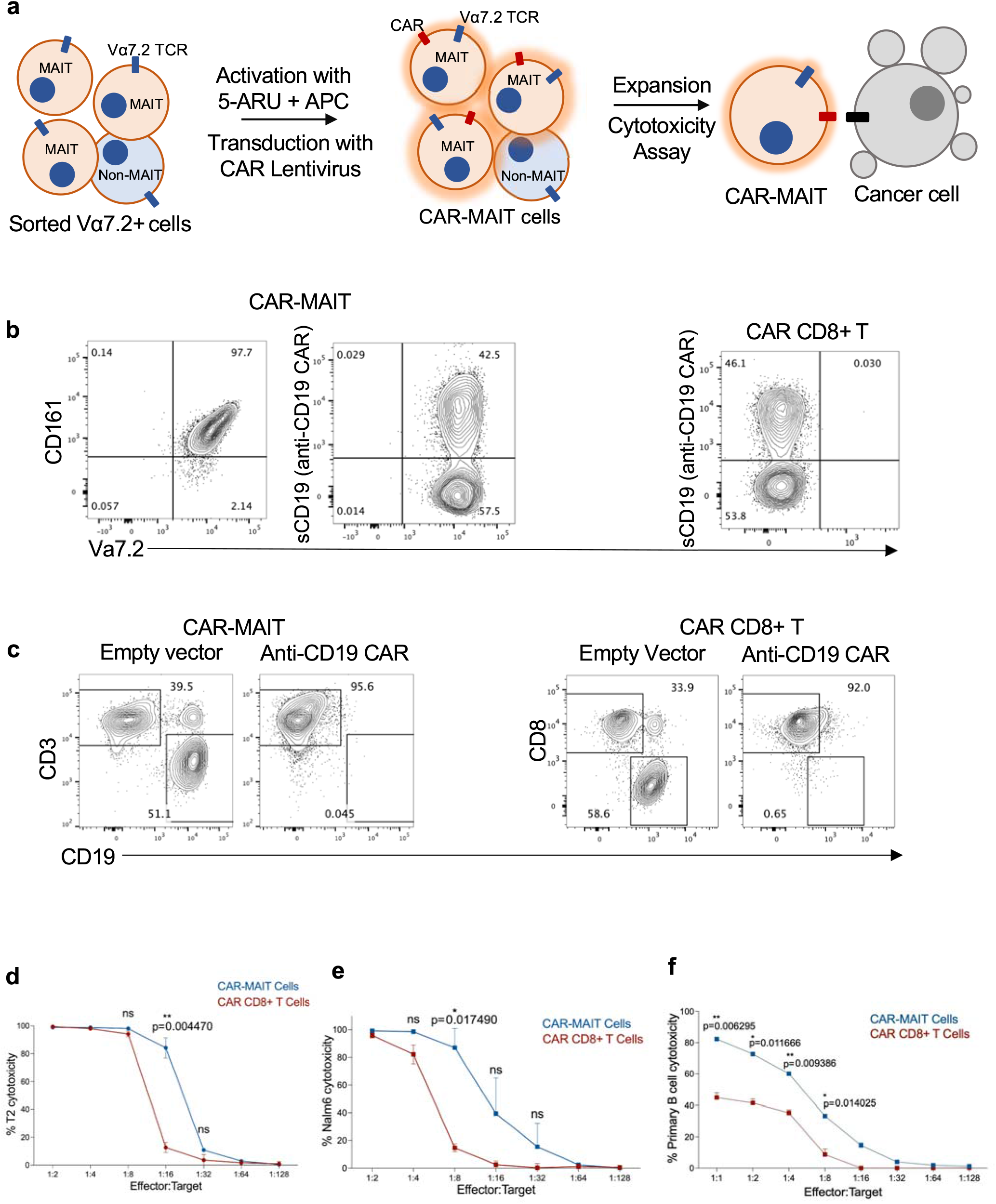
Development and generation of purified CAR-MAIT cells and cytotoxicity towards B cell lymphomas. **a**. Schematic illustration demonstrating the sorting, engineering, and cytotoxicity assay strategy for CAR-MAITs. Vα7.2+ cells were isolated from CD4 negative (CD4-) PBMC population. Blue rectangles on the cell surface represent Vα7.2 T cell receptors. The cells with orange and blue cytoplasm show MAIT and non-MAIT cells, respectively. Sorted Vα7.2+ cells were activated by adding the 5-ARU molecule in the presence of autologous MR1-expressing Antigen Presenting Cells (APCs). Red glow around the cells represent activation. Selectively activated MAIT cells were then transduced with CAR encoding lentiviruses and expanded in IL-2 media. Red rectangles on the cell surface represent chimeric antigen receptors. Finally, CAR-engineered MAIT cells were combined with target cells for cytotoxicity assays, black rectangle on the cell surface represents CD 19 protein, b. Flow cytometry plots demonstrating CAR surface expression in CAR-MAITs and CAR CD8+ T cells. Engineered CAR-T cells were stained with CD19-Fc fusion protein then with an anti-Fc antibody to determine the expression levels of anti-CD19 CARs. **c**. Representative flow cytometry plots of cytotoxicity assays in which CAR-MAITs and CAR CD8+ T cells were combined with T2 cells at an effector:target cell ratio of 1:1 and 1:2, respectively. The effector to target cell ratio was calculated based on the number of effector cells expressing CARs. CAR-MAIT cells, CAR CD8+ T cells and the target cells were identified by the expression of CD3, CD8 and CD19, respectively. Empty vector transduced MAIT and CD8+ T cells were used as a control, **d, e**. Line graphs of killing percentages from the cytotoxicity assays of CAR-MAITs and CAR CD8+ T cells targeting T2 and Nalm6 cells. Red and blue lines represent CAR CD8+ T and CAR-MAIT cells, respectively. The effector to target ratio was titrated from 1:2 to 1:128 with 2-fold reciprocal dilutions of effector cells. Percent cytotoxicity was calculated based on the percentage of cell death in experimental conditions in relation to control conditions, f. Line graph showing the percent cytotoxicity of CAR-MAIT and CAR CD8+ T cells on primary B cells at ratios ranging from 1:2 to 1:128. Cytotoxicity assays were replicated twice with different donors. Error bars represent one standard deviation of mean value. Two-tailed unpaired t tests were used to determine the statistical significances. The number of replicates in figures **c** and **d** were 3 and 2 for figure **e**.

**Figure 3:**
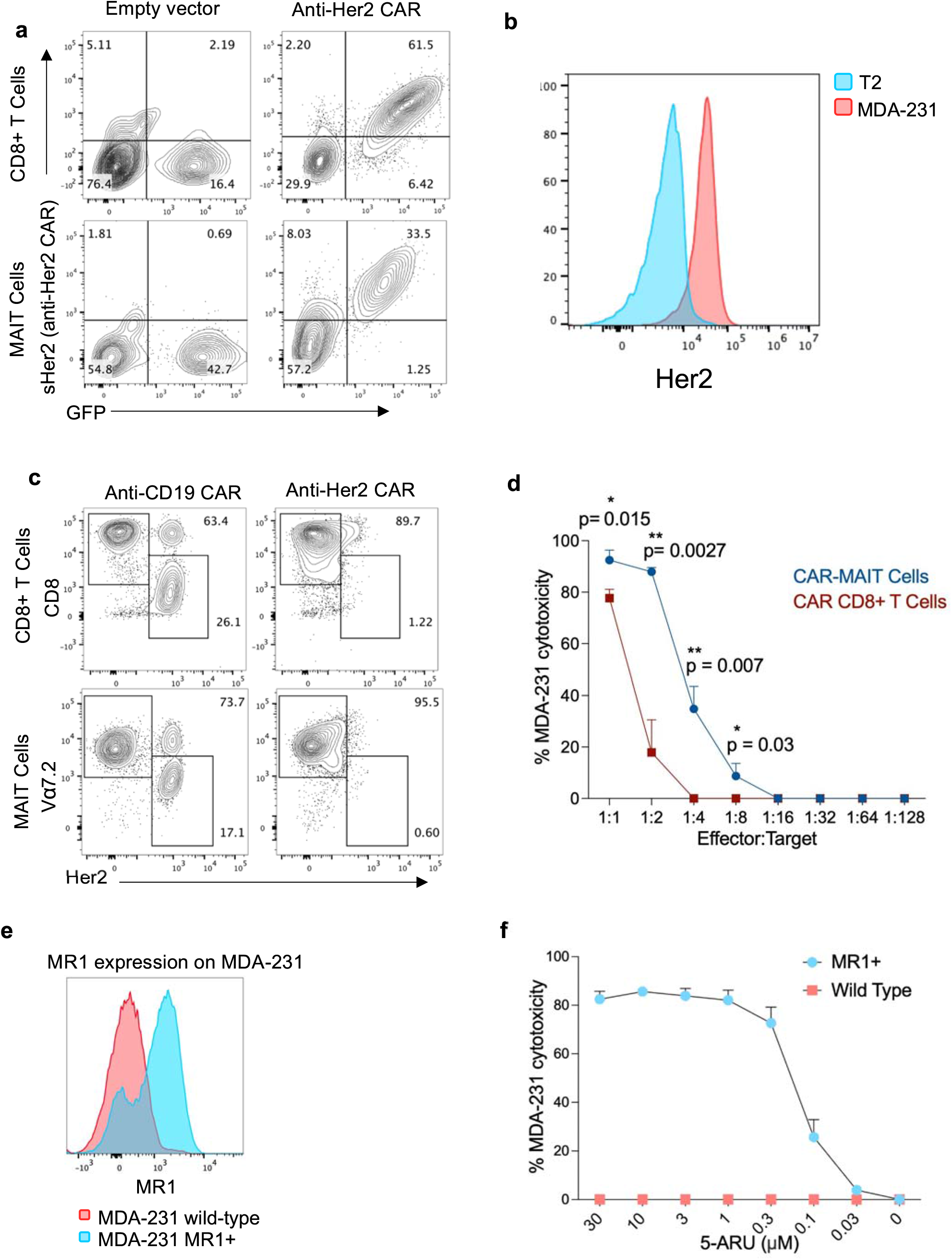
Cytotoxicity of MAIT cells against a solid tumor cell line: MDA-231. a. Flow cytometry plots demonstrating the anti-Her2 CAR surface expression in CAR-MAITs and CAR CD8+ T cells. Engineered CAR-T cells were stained with Her2-Fc fusion protein and an anti-Fc antibody to demonstrate the expression levels of anti-Her2 CARs. GFP lentivirus markers were also used to show the co-expression with anti-Her2 CAR in engineered T cells, b. Flow cytometry overlays of MDA-231 cells stained with an anti-Her antibody. The red population in the histogram represents T2 cells as control whereas the blue population shows the presence of Her2 expression in anti-Her2 stained MDA-231s. c. Representative flow cytometry plots of cytotoxicity assay using CAR-MAITs and CAR CD8+ T as effector cells and Her2+ MDA-231 as target cells. MAIT cells, CD8^+^ T cells and MDA-231 cells were identified with CD8, Vα7.2 and Her2 expression, respectively. Effector (CAR expressing T cells) to target ratio of 1:1 is shown, d. Line graph showing the percent cytotoxicity of CAR-MAITs and CAR CD8+ T cells targeting MDA-231 cells. Red and blue lines represent CAR CD8+ T and CAR-MAIT cells, respectively. Error bars represent one standard deviation of mean value. The effector to target cell ratio was calculated based on the number of effector cells expressing CARs. The effector to target ratio was titrated from 1:2 to 1:128 by 2-fold reciprocal dilutions of effector cells. Percent cytotoxicity was calculated based on the percentage of cell death in experimental conditions in relation to control conditions. Two-tailed unpaired t test was used to determine the statistical significances, e. Histogram overlay demonstrating the MR1 expression levels in wild-type MDA-231 cells and MR1 overexpressing MDA-231 cells. Red and blue populations indicate wild type and MR1 overexpressing MDA-231 cells, respectively, f. Non-infected MAIT cell cytotoxicity towards wild-type and MR1 overexpressing MDA-231 cells in the presence of different concentrations of 5-ARU. MAIT cells were purified, activated and expanded as described in the Methods section. After 3 weeks of expansion in IL-2 containing media, MAIT cells were cultured with wild-type (red squares) and MR1-overexpressing (blue circles) MDA-231 cells in the presence of 5-ARU. 5-ARU was titrated from 30 μM to 0.003 μM by 3-fold reciprocal dilutions. Cytotoxicity assays were replicated twice with different donors.

We next sought to compare the cytotoxic function of CAR-MAITs and conventional CAR CD8+ T cells. For this, we generated CD8+ anti-CD19 CAR-T cells by lentivirus transduction upon CD3/CD28 activation. Both CAR-T cell subsets were then expanded for 10-12 days in IL-2 containing media. Staining for anti-CD19 CAR surface expression revealed that both cell types expressed CD19-CAR at comparable levels (Fig. 2b). The effector cells were then co-cultured for 3 days with T2, Nalm6 or primary B cells, all of which express the target antigen CD19 on their surface (Supplementary Fig. 1b, c). Both conventional CAR CD8+ T and CAR-MAIT cells almost completely killed the target cells, while no cytotoxicity was observed by cells expressing empty vector controls (Fig. 2c), at different effector to target cell ratios (Fig. 2d, e). Interestingly, CAR-MAIT cells displayed slightly more cytotoxicity to T2 (p=0.0044) and Nalm6 (p=0.0174) than conventional CAR CD8+ T cells did at the effector/target cell ratio of 1:16 and 1:8, respectively (Fig. 2d, e). CAR-MAIT cells had more cytotoxicity also towards primary B cells compared to CAR CD8 T cells (Fig 2f). A representative flow cytometry data plot of a primary B cell cytotoxicity assay is shown in Supplementary Figure 2.

### Cytotoxicity of CAR-MAIT cells against a breast cancer cell line

We next engineered anti-Her2 CAR-MAIT cells to test their efficiency against Her2 expressing cells, as some solid tumors have high Her2 expression(22). The anti-Her2 CAR cassette was constructed by switching the extracellular domain of anti-CD19 CAR with an anti-Her2 single chain variable fragment (scFv) as described in the Methods. Anti-Her2 CAR expressing conventional CD8+ T cells were also generated to compare the cytotoxic features of both effector cells. Surface staining of anti-Her2 on CAR-MAIT and CAR CD8+ T cells revealed that both engineered effector cell types expressed comparable levels of CAR, which correlated with the GFP reporter expression (Fig. 3a). Breast cancer cell line MDA-231 cells were stained with an anti-Her2 antibody to confirm its expression (Fig. 3b). We next determined the cytotoxicity of these CAR-engineered cells to the MDA-231 cells. Effector cells expressing anti-CD19 CAR were used as negative controls. Both anti-Her2 CAR-MAIT and anti-Her2 CAR CD8+ T cells efficiently killed MDA-231 cells (Fig. 3c) and CAR-MAIT cells showed significantly higher cytotoxicity than conventional CAR-T cells at the effector/target cell ratios of 1:1 (p=0.0015), 1:2 (p=0.0027), 1:4 (p=0.007) and 1:8 (p=0.03) (Fig. 3d).

We also tested the cytotoxicity of non-infected primary MAIT cells against MR1 expressing cancer cells in the presence of 5-ARU. For this, we engineered MDA-231 cell lines to overexpress MR1, then used them as target cells in a cytotoxicity assay with primary MAIT cells in the presence of 5-ARU. Engineered MDA-231 cells were stained with an anti-MR1 antibody and MR1 overexpression was confirmed (Fig. 3e). Primary MAIT cells were purified, activated and expanded as described in methods without adding any lentivirus. After 3 weeks of expansion in IL-2 containing media, non-infected MAIT cells were cocultured with wild-type or MR1 overexpressing-MDA-231 cells in the presence of different concentrations of 5-ARU for 2 days. Flow cytometry analysis revealed that primary MAIT cells were able to kill MDA-231 cells overexpressing MR1 in in the presence of different concentrations of 5-ARU, in a dose dependent manner, whereas no cell death was observed in the wild-type MDA-231 condition at any concentration (Fig. 3f and Supplementary Figure 3a). CD25 staining showed activation of MAIT cells in cocultures treated with 5-ARU (Supplementary Figure 3b).

### Cytokine secretion by activated CAR-MAIT and Conventional CAR-T cells

We next determined the cytokine response of CAR-MAIT cells upon triggering of CAR signaling. A multi-plex flow-based immunoassay was used to measure TNF-γ, TNF, Granzyme B (GzB) and Granulocyte-macrophage colony-stimulating factor (GM-CSF) levels in the supernatants of effector/target cell co-cultures at a ratio of 1:1. CAR-MAIT cells secreted significantly lower levels of TNF-γ than CAR CD8+ T cells in cytotoxicity assays targeting primary B cells (p=0.003), T2 cells (p=0.006), Nalm6 cells (p=0.002) and MDA-231 cells (p<0.0001) (Fig. 4a). However, TNF secretion by CAR-MAIT cells was significantly higher compared to CAR CD8+ T cells against primary B (p=0.0009), T2 (p=0.0423) whereas lower against MDA-231 cells (p=0.0079) (Fig. 4b). We did not observe a significant difference in TNF expression between the two effector cell types when targeting Nalm6 cells (Fig. 4b). Similar to TNF-γ, GzB and GM-CSF levels were also significantly lower in CAR-MAIT cytotoxicity conditions targeting primary B (p=0.0023 and 0.0003, respectively), T2 (p=0.002 and 0.0009, respectively), Nalm6 (p=0.0011 and 0.0005, respectively) and MDA-231 cells (p=0.0367 and 0.0002, respectively) (Fig. 4c, d).

**Figure 4:**
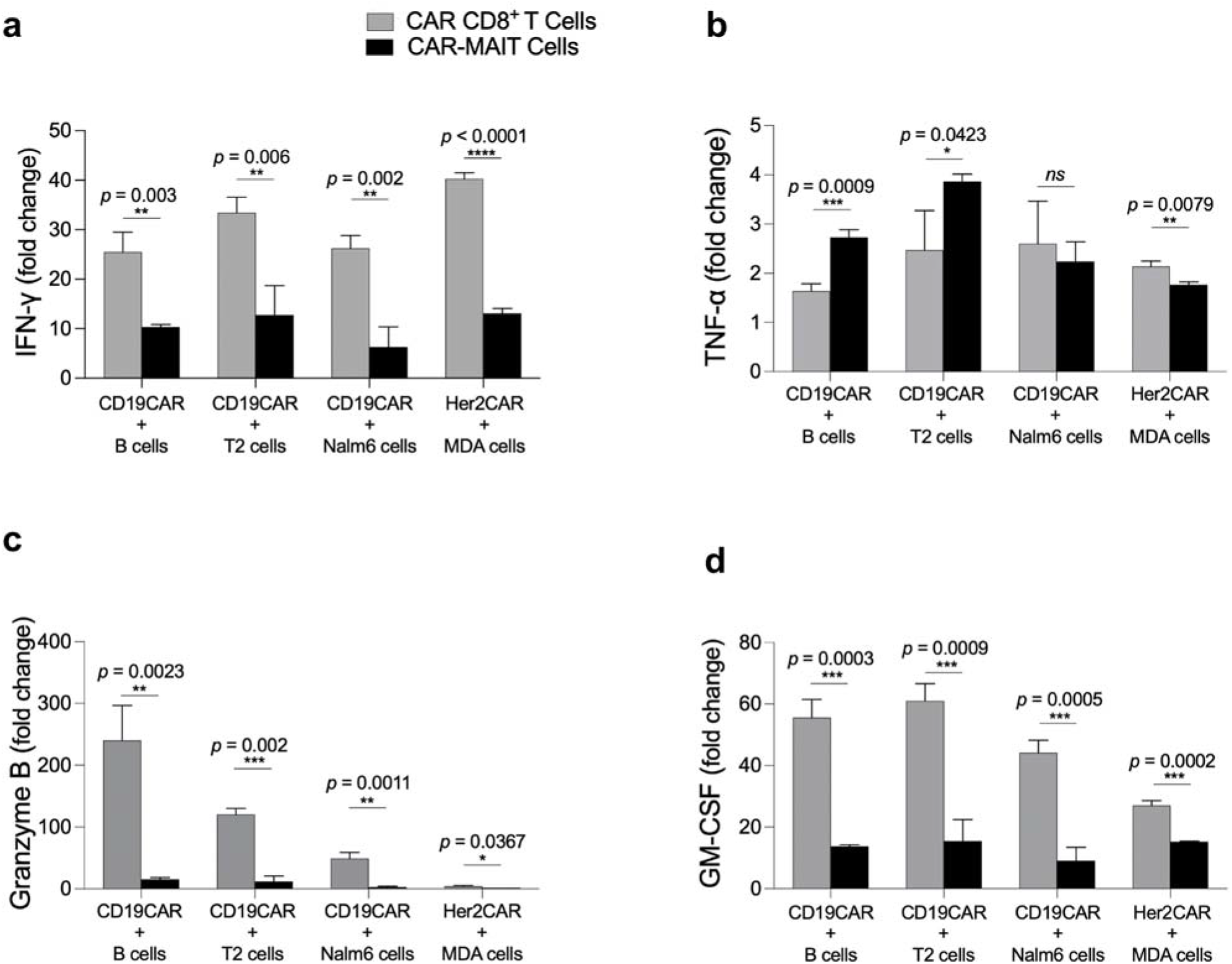
Cytokine secretion of CAR CD8+ T and CAR-MAIT cells, upon activation. In the supernatants of co-cultured CAR-MAIT and CAR CD8+ T effector cells with primary B, T2, Nalm6 or MDA-231 target cells, shown in gray and black bars, respectively, following cytokines were measured and expressed as fold change: a. TNF-γ, b. TNF, c. Granzyme B (GzB) and d. GM-CSF. Fold changes were calculated based on the fold increase of experimental conditions compared to control conditions. Cytotoxicity assays were replicated twice with different donors. Error bars represent one standard deviation of mean value. Two-tailed unpaired t test was used to determine the statistical significances.

## Discussion

Cancer immunotherapies exploiting chimeric antigen receptors show great promise in treating certain tumor types. However, there are major disadvantages such as a high risk of adverse effects like cytokine release syndrome, neurotoxicity, anaphylaxis and graft versus host disease, lower efficacies against solid tumors compared to hematological malignancies and also practical issues such as the need to utilize the patient’s own T cells for each treatment, which increases the cost and prevents off-the shelf treatment options.(23, 24) In order to overcome some of these hurdles, we sought to develop a novel CAR-T cell with a non-classical T cell subset, MAIT cells, against a variety of tumor types. Here we demonstrate highly efficient generation of CAR-MAIT cells, which display equivalent or higher cytotoxicity to target cells, compared to conventional CAR-T cells.

Conventional T cells recognize antigens in the context of major histocompatibility complex-1 (MHC-I) or class-11 (MHC-II) molecules on the presenting cells, which results in an allogeneic reaction if the cells derived from non-self-donors are used for treatment. Thus, current CAR therapies utilize the patient’s own T cells that are separated by leukapheresis, engineered *in vitro* and then transferred back to the same patient. This limitation to use only autologous cells prevents development of off-the-shelf cell-based therapies, which would greatly improve the process, making it a cheaper, faster, and more readily available treatment option.

Recently, Invariant Natural Killer T (iNKT) cells are being used in the adaptive immunotherapies to address this issue.(25-28) iNKT cells are also increasingly studied for their distinct features, including their robust anti-tumor activity, their intrinsic nature of regulating other immune system members, and their susceptibility to engineering; namely the qualities they share with MAIT cells (29). Following their activation with αGalSer, iNKT cells were used in preclinical studies on patients with multiple myeloma, adenocarcinoma, renal cell carcinoma, non-small-cell lung cancer, and large-cell carcinoma with encouraging results (1). Similar to iNKT cells, MAIT cells also do not employ the MHC-I molecule for antigen recognition and conceivably lack the allogeneic adverse reactions. The advantage of MAIT cells to iNKT cells is the abundance of MAIT cells in the human immune system compared to low frequencies of iNKT cells.(30) In this regard, MAIT cells have the potential to be the main cell type in the future of adaptive off-the-shelf cancer immunotherapies.

To be able to generate engineered human MAIT cells and proliferate them to a degree at which they can be employed against target cells, we optimized an expansion protocol. The protocol we designed in our study for the MAIT cells using 5-ARU, similar to using αGalSer for iNKT cells, is of great importance in the field of cell-based immunotherapy since it allows us to engineer and make use of this distinct and abundant human T cell subtype. To our knowledge, this is the first study to demonstrate the feasibility of using chimeric antigen receptor engineered MAIT cells against specific targets.

Our key finding was the demonstration that MAIT cells engineered with chimeric antigen receptors perform cytotoxicity against specific cancer targets. This finding encouraged us to perform extensive cytotoxicity assays comparing CAR-MAITs with conventional CAR CD8+ T cells. Remarkably, we found that CAR-MAITs show similar or, in some cases, significantly higher cytotoxicity compared to the conventional CAR CD8+ T cells in our in-vitro assays. In future studies, it will be important to address the efficacy and potential side effects of CAR-MAIT cells in in-vivo experiments. For example, how specifically CAR-MAIT cells target cancer cells compared to normal tissue expressing similar antigens? How long will they persist in vivo? Will they be effective in solid tumor microenvironments?

The second key finding in our study was that despite robust cytotoxic capacity, CAR-MAIT cells secrete significantly less TNF-γ, GzB and GM-CSF compared to conventional CAR-T cells. Indeed, a major pitfall in cell mediated adaptive immunotherapies is the severe adverse effects of the over T cell activation, which results in high pro-inflammatory cytokine secretion(31). This in turn can lead to cytokine release syndrome and neurological toxicities, which are two of the biggest concerns after the infusion of engineered CAR T cells(32). Thus, lower cytokine secretion and high cytotoxic activity of CAR-MAIT cells compared to conventional CAR CD8+ T cells could be of importance in developing safer and more effective immunotherapeutic effector cells. In fact, reducing GM-CSF has been demonstrated to decrease cytokine release syndrome and neuroinflammation but enhance CAR-T cell function.(33) Reduced TNF-γ levels could mean CAR-MAITs may be less potent at stimulating endogenous anti-tumor response(34) and lead to worse prognosis or may be beneficial by promoting less inflammation in the tumor microenvironment compared to conventional CAR CD8+ T cells. These are also some of the questions that need to be addressed via in-vivo studies comparing conventional and MAIT CAR-Ts. Together, these findings could be particularly relevant for adaptive immunotherapies using lower doses of CAR-MAITs compared to CD8+ CAR-T cells, which may result in better therapeutical efficiencies while minimizing adverse effects to normal cells or tissues.

Another key outcome in our study was engineering CAR-MAITs to recognize and kill target cells which express Her2 on their surface more efficiently than conventional CAR CD8+ T cells. This result was critical since Her2 is one of the surface molecules highly expressed on several solid tumors(22). Additionally, it has been previously reported that primary MAIT cells can infiltrate tissue sites of chronic inflammation(35) and are abundant in the mucosal-associated and solid tumor microenvironment(16-18). Therefore, this migration capacity towards sites of inflammation can be of advantage in employing CAR-MAIT cells for solid tumor targeting immunotherapies. It is also reported that MAIT cell accumulation in tumor microenvironment is associated with poor survival in colorectal cancer patients(36). Engineering MAIT cells with CARs to recognize tumor cells in solid tissues, can turn this caveat of MAIT cell infiltration into tumor microenvironment to an advantage in a cell-based therapy approach described herein.

In addition, we also showed that primary MAIT cells were able to kill MR1 overexpressing target cells in the presence of 5-ARU in a dose dependent manner. Because MR1 can be expressed by several tumor types and that MAIT cells are found to be increased in some cancer niches such as primary colorectal lesions, hepatic metastatic lesions or multiple myeloma(37) it may be possible to exploit their 5-ARU-mediated activation to unleash their anti-tumor activity. It is also conceivable to combine both MR1-restricted and specific CAR engineering to fine tune MAIT anti-tumor responses, especially in more challenging solid tumor microenvironments. It will be important in future studies to further develop these engineering approaches and test them in vitro in 3D-models of cancer and ensure that these cells can effectively infiltrate tumor environment and endowed with fail-safe mechanisms.

Future animal experiments will help to observe the in-vivo behaviors of the CAR-MAIT T cells in detail and will shed light on this approach from different perspectives.

Development of CAR-MAIT cells and other innate-like T cells with unique feature sets can accelerate novel adaptive immune cell-based approaches in cancer immunotherapy and overcome some of the key challenges in the quest to utilize immune system as an important cancer treatment.

## Data availability

The source data for the Figures along with the Supplementary Figures are available upon request.

## Acknowledgements

We thank Sarah Cassidy and Courtney L. Gunter for careful review and editing of the manuscript.

## Author Contributions

M.D., E.K., L.K. and D.U. designed the experiments, M.D., E.K., L.K., L.P., X.C., M.Y. carried out the experiments. M.D., E.K., L.K., D.U. analyzed the data. M.D. and D.U. wrote the manuscript.

---

**Supplementary Figure 1:**
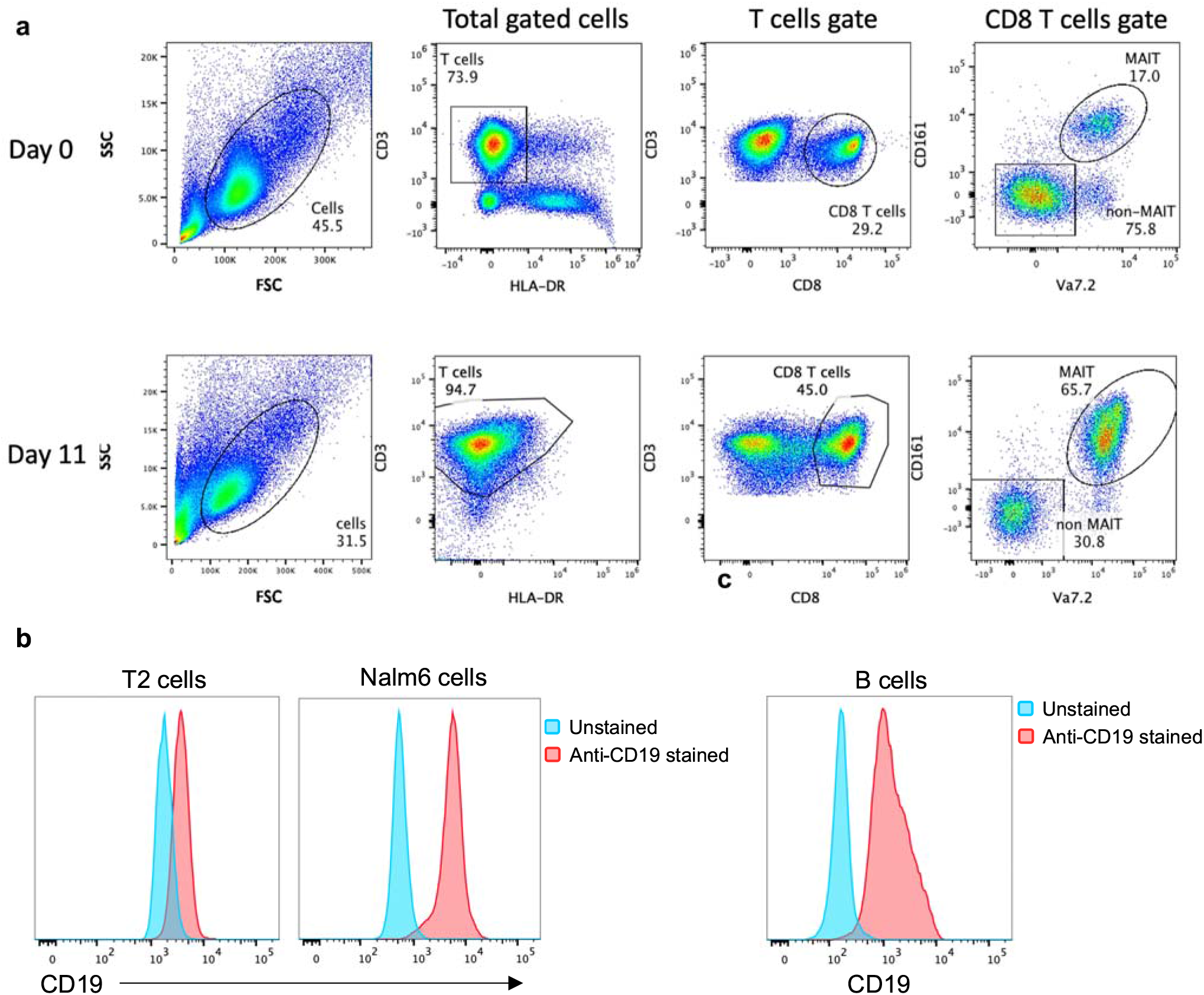
Gating strategy for CAR-MAIT cell expansion and CD19 surface expression of Nalm6 and T2 cells. **a**. Flow cytometry plots showing the gating strategy of CAR-MAIT expansion experiments at days 0 and 11. **b**. Overlay plot of flow cytometry data of T2 and Nalm6 cells stained with an anti-CD19 antibody. Blue and red populations demonstrate unstained and anti-CD19 antibody-stained cells, respectively, **c**. CD19 expression of primary B cells.

**Supplementary Figure 2:**
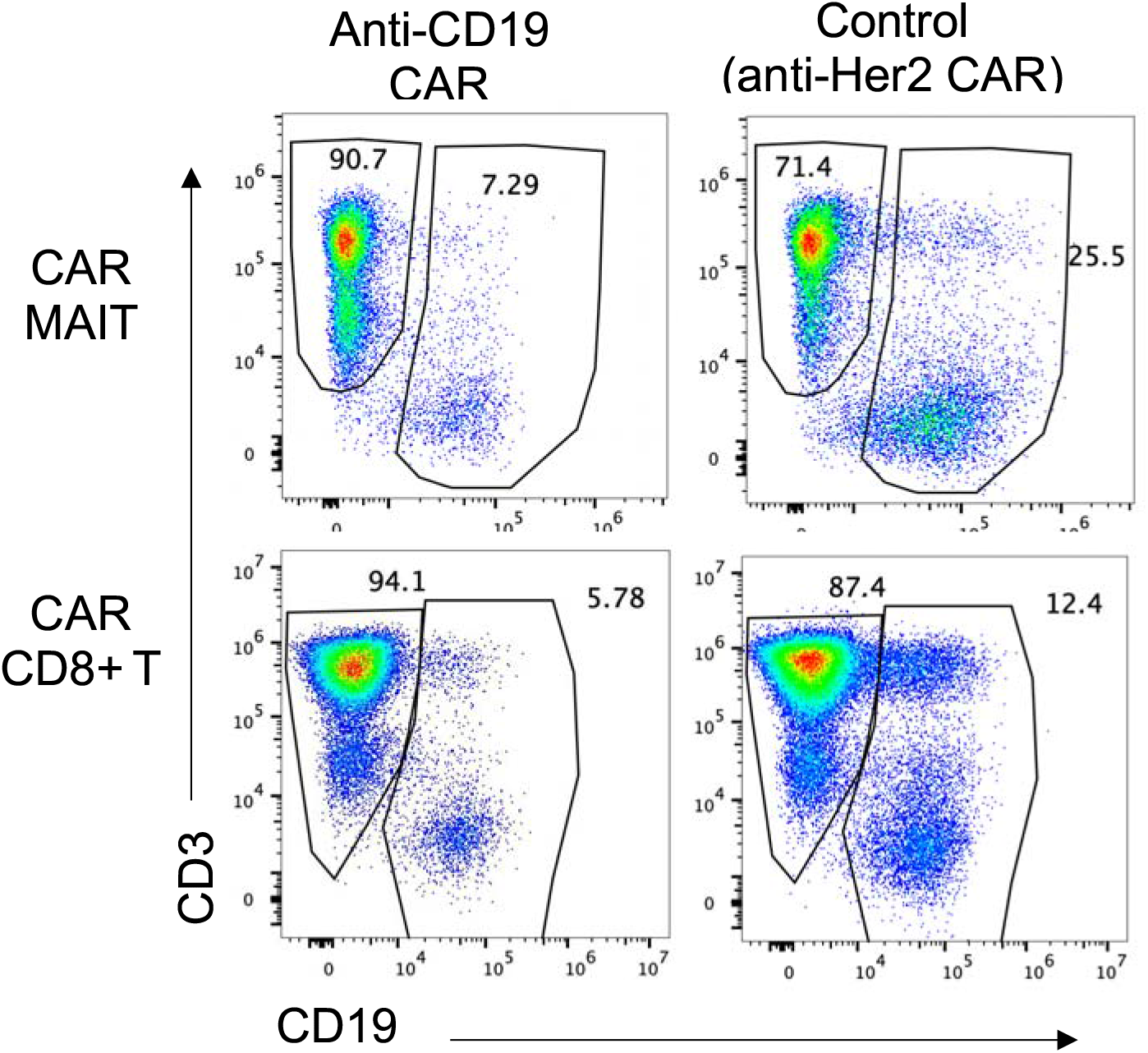
Cytotoxicity of CAR-T cells towards primary B cells. Representative flow cytometry plot of B cell cytotoxicity assay. Anti-Her2 CAR expressing CAR-MAIT and CAR CD8+ T cells were used as controls.

**Supplementary Figure 3:**
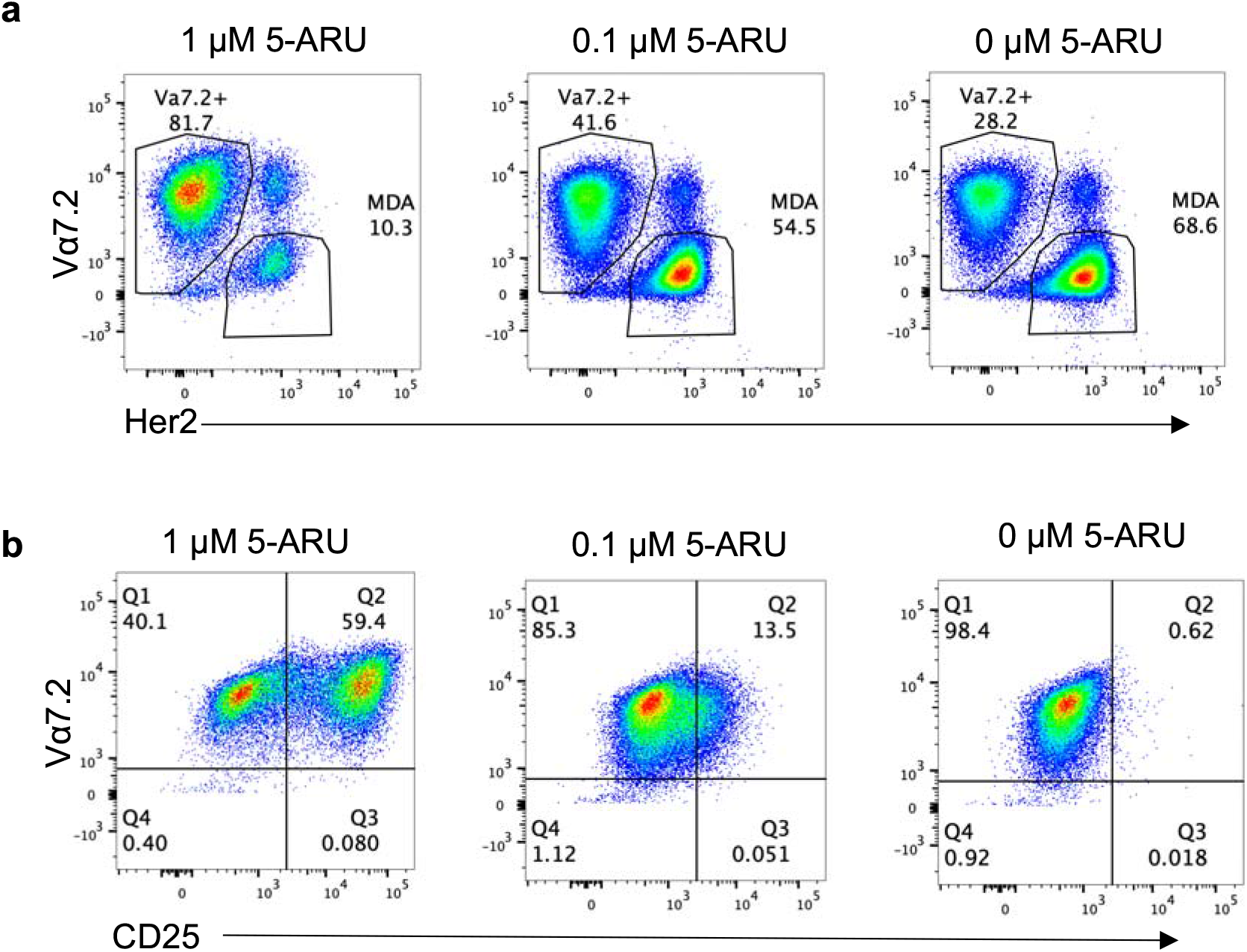
Primary MAIT cell cytotoxicity to MR1 expressing MDA-231 cells in the presence of 5-ARU. Representative flow cytometry plots of MAIT cell cytotoxicity to MR1 overexpressing MDA-231 cells in the presence of different concentrations of 5-ARU. **a**. Percentage of cells are shown in the cytotoxicity assay. Primary MAIT cells and MDA-231 cells were identified via their Vα7.2 and Her2 expressions, respectively, **b**. Flow cytometry plots demonstrating the activation of primary MAIT cells during the cytotoxicity assay described in **a** in the presence of 5-ARU. CD25 expression was used to determine the activation.

**Supplementary Figure 4:**
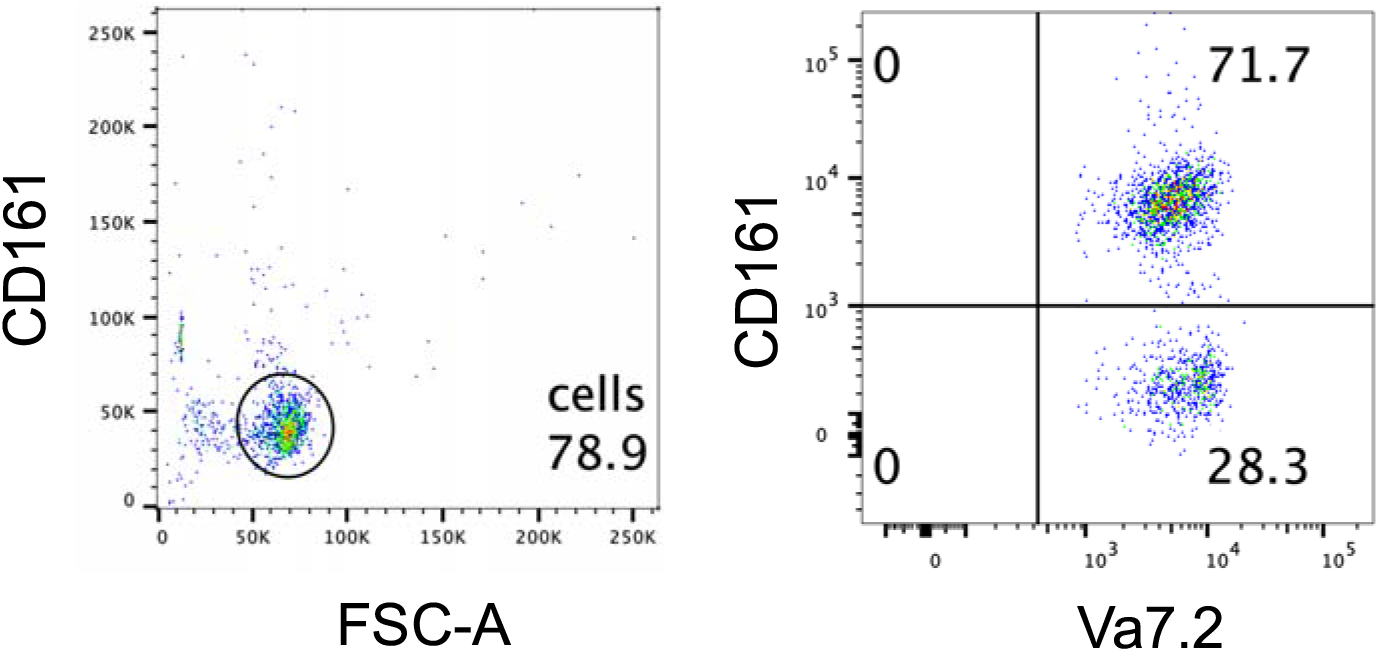
Post-sort purity of MAIT cells. Representative flow cytometry plot of MAIT cell purity following sorting. CD161 and Vα7.2 were used as MAIT cell markers.

